# Trends of the Passerines avifauna are linked climatic variability and habitat management in a protected artificial wetland in southeast Iberia

**DOI:** 10.1101/2025.10.21.683676

**Authors:** Ignacio García Peiró

**Author notes:** Corresponding author: SEO/Birdlife-ALICANTE. C/ El Salvador, 17-4D. 03203 Elche (Alicante). ES-Spain.

## Abstract

The Mediterranean region is facing significant impacts on populations of biota, particularly birds, due to climate variability and habitat alteration. Consequently, the ecological diversity of many ecosystems has diminished, and the abundance of bird species has declined. This study employs a novel strategy in a seasonal freshwater artificial wetland in south-east Iberia establishing species-specific index trends for common passerine birds by comparing them with climate and habitat fragmentation data. A climatic overview indicates minimal trends towards warmer and wetter conditions in terms of temperature and rainfall over the 15-year study period. This pattern persisted over a 30-year period, resulting in the formation of a Mediterranean climatic sub regional scenario with warm, dry conditions that may influence the passerine assemblage in an analogous manner to that observed in other Mediterranean assemblages. One notable feature is that this assemblage in the 1990s was characterized by low natural diversity and specialization when it was a highly disturbed wetland. However, it became natural by the mid-2000s, with diversity values like those observed in other central Mediterranean wetlands. Trends used to evaluate species-specific patterns revealed that 98% of species had an unknown trend status during the study period, with a trend slope ranging from 0 to 15%, which was not substantially different than zero, probably due to the unequal standardization methodology and the short time studied. Further research using Generalized Linear Mixed Models (GLMM) revealed that precipitation and habitat alteration had the most detrimental effect on species. Of the observed species, 60% were affected by both factors, the majority of which were Palearctic (20%). The reduction of wintering areas and shifting migration routes may account for these effects. Further research is required on longer temporal data to clarify the evolution of the avifauna in this wetland.

## 1. Introduction

The habitat and space provided by the Mediterranean are very favorable for some of the North African faunal traits that are exclusive to the southern Mediterranean, especially in the south-part of Iberia (Cheylan 1991, Martínez-Freiría et al. 2010, Husemann et al. 2014). In the class Aves, passerines represent a real hotspot of biodiversity (Covas & Blondel 1998). One of its most interesting features is the scarcity of Mediterranean bird species, with European and Palearctic elements making up the majority of the region’s composition (Voous 1960, Blondel 1987). They are important elements of study of the evolutionary processes (Pons et al 2016).

Freshwater areas, particularly wetlands (marshes, lagoons, swamps), are declining in number and surface area due to significant water deficits caused by climate change since the turn of the century (Perennou et al. 2012; Lefebvre et al. 2019). The loss of Mediterranean wetlands has reduced the biological diversity of passerines (Paracuellos 1993, Peiró 2018), many of which are extremely important European-Turkestanian or Turkestanian-Mediterranean faunistic elements (Kirwan et al. 2010, Rustamov 2015, Peiró & Esteve 2002). These elements are vital for biodiversity conservation, ecological study, and understanding evolutionary processes since they play an important role in the early stages of plant successions (see Schulze-Hagen and Leisler 2011, Abramova 2019) and as secondary habitat (Bozó et al 2020).

The preceding assumptions suggest that the loss of specialists may enhance the extinction risk of ecosystems through biotic homogenization, with generalist winners replacing losers’ specialists (McKinney & Lockwood 1999; Le Viol et al. 2012). This is primarily due to anthropogenic causes (Smart et al. 2006, De Victor et al. 2008a), while climate can play an influence (Davey et al. 2011).

The shift towards patchy wetlands (Paracuellos 2006 & 2008) has facilitated the proliferation of noteworthy Palearctic specialist songbirds across Europe, which congregate in reed-bed masses. This phenomenon is exemplified by the Bearded Reedling *Panurus biarmicus* (Erard 1966, Kumberloeve 1968, Mead & Pearson 1974, Loison & Godin 1975, Olsson 1975, Antoniazza & Levéque 1977, Sluys 1982) is listed as Vulnerable in Spain (Vera 2021). In the study area, their small populations are situated at the southernmost extent of the species’ distribution in Western Europe (Gosler & Mogyorósi, 1997) and may have dispersed to southern Iberian counterparts from the south-east of France, where this species was equally common and highly specialized since 1830, and where temperatures increased from 1960 (Galewski & Devictor, 2016). Currently, the species is almost extinct in the study area (Belenguer et al., 2016).

Inside this scenario, the analysis of bird trends using different monitoring data has been highly important to detect the impact of climate change on songbird trends, and applied in Western Europa (Reif et al. 2008, van Turnhout et al. 2008) and Eastern Europe (Zalakevičius 1998, Bozó et al. 2023, Tamás et al. 2023), and the results have shown to be highly variable (Jiguet et al. 2010), but few studies yet don’t make use the TRIM methodology despite Bird Monitoring Schemes were set long ago (Vorisek & Marchant 2003, Vorirsek et al. 2008). However, it appears that little trend research has been carried out in wetlands (Žalakevičius & Švažas 2005, Van Turnhout et al. 2010) and many few using long-term ringing data and showing an overall decrease in the European avifauna (Petras & Vrezec 2022).

Based on the aforementioned arguments, the current study aims to: 1) provide a comprehensive overview of the climatology and zoogeography of the study area that has not been yet adequately covered (Peiró & Esteve 2001, Peiró 2006) and 2) address the question of how much abiotic (local climate) and biotic (anthropogenic) factors can influence the population trends of a Mediterranean wetland passerine assemblage subject to significant artificial management of water cycles and vegetation, as well as how these factors affected the homogenization of the assemblage and provide additional insights for the years to come.

## 2. Material and Methods

### 2.1. The study region, area and site

With an estimated surface area of 260,000 Ha, the extreme southeastern Iberian span the easternmost portion of Iberia’s Baetic southern plateau (Armada et al. 2018). Almería (37°58’N - 2°45’W), Murcia (37°58’N - 01°33’W), and Alicante (38°34’ N - 00°48’W) are its three main provinces. The region has a semiarid Valencian climate (Costa 2019) and a Thermo-Mediterranean Iberian climate (Rivas-Martínez 1983, Rivas-Martinez & Armaiz 1984). The weather is seasonal, especially in the spring and autumn when it’s rainier.

The study area “*El Hondo Natural Park*” (abbreviated EH, 38°’16’N - 00° 41’W) is in the middle-south of the province of Alicante. It is a 2.400-hectare inland freshwater swamp reservoir that was completed around the turn of the 20th century (Medina 1985) and designated as a regional natural park in 1988 (GVA 1988). Additionally, it was referred to as “El Hondo Swamp” in 1989 as RAMSAR site number 14 (Potenciano & Andueza 2006) and renamed as site number 455 (Bernués & Mateache 2020) because it maintains large populations of endangered waterbirds (Medina & Robledano 1995). A rich community of medium-to-small reed-bird species, some of which are currently threatened, and developed inside the eutrophic vegetation between man-made structures (Medina & García 1984, Peiró 1999).

The 800-meter line between two ponds is sited in the core area of the park and forms a succession of halophytic vegetation that reaches the park’s boundaries (Peiró & Esteve 2001). The line included of saltmarshes of the species *Salicornia*, *Suaeda*, and *Arthrocnemum*, as well as common reeds *Phragmites australis* at various growth stages (Navarro 1985, Peiró 2006).

### 2.2. The Climatic data treatment

I used annual mean temperatures and rainfall as indexes of abiotic factors or climatic variability. Forecast variables were collected from the “*Estación Experimental Agraria of Elche (EEA-IVIA)*”, a meteorological station located 11 km northeast of the park at 38°14’N 1°25’W and 62 m above sea level. The data was gathered over a 15-year period (1991-2005, period D1) in EEA notepads and online from IVIA http://riegos.ivia.es/datos-meteorologicos. Long-time datasets for a 30-year period (1991-2020, period D2) were used in comparison with the first dataset. The climatic data consisted of 1) annual and monthly average temperatures (°C) and 2) overall annual and monthly precipitation (mm). To see the overall summary of climate, additional variables were used: 1) mean annual temperature in spring (March-May), summer (June-September), autumn (July-October), and winter (November-February); and 2) overall annual winter (November-February).

The research area’s bird seasonality is achieved, however certain climatic data, particularly the annual temperature did not reach normality (Wilk’s Shapiro test: W = 0.815, P = 0.006). However, because rainfall varies so much, parametric tests based on the alternative hypothesis of outlier existence were employed (Grubbs Test, Grubbs 1969). Standard deviations (SD) were shown with the means. Two-tailed significance values were set at P < 0.05.

### 2.3. The Monitoring management

Strong management practices were implemented at the research site, mostly using human and animal grazing and harvesting (wildfires, reed cutting, and weed-clearing base) (Peiró 2006). By doing these steps, we were able to raise the reed’s height and gradually remove its structure during the study period. The yearly average height of reeds was selected as a measure of habitat change brought about by management. From 1991 to 2005, I measured the height of the reed beds on both sides of the study path in order to collect field data for assessing these changes. One to four months following each management intervention, I randomly picked one to twenty reed stems and used a meter rule to measure the height (in cm) from the base to the seeds. Management actions were conducted almost every year during the study period and consisted mainly in harvesting, grazing and wildfires of the reedbed path.

### 2.4. The Bird’s data recording and monitoring

First, throughout annual surveys (1991-2005, N = 15, period D1), the author implemented a Flexible Constant Effort Site mist-netting program in the Park’s core region with varied ringing visits, which typically lasted four hours after sunrise. Only comparisons of biodiversity were conducted using the incomplete datasets from 2006 to 2012 (N = 7, period D3). According to Peterson et al. (1977) and Svensson (1992), I metal-ringed every bird (Ministery of Environment-MADRID, SPAIN), measured each bird, and then released them all to the same location. Nearly every season from D1 was covered by the ringing and handling in D1 and D3 (1) (N = 15 years). I attempted to evaluate potential assemblage changes over long periods of time using some standardized ringing data from period D3.

I mostly used period 1 for data monitoring. By catching them, ringed birds were measured and standardized. Standardization procedures consisted by dividing the number of individuals by trapping effort [(individuals/(m2·hour)) ·1000]. This was done differently for each of the following annual presences at the site: breeding (March-October, B), wintering (November-February; W), sedentary (January-December, S), spring migrant (March-May, SM), autumn migrant (July-October, AM), and occasional (O) (Table 1). Sum spring-autumn trapping effort (SO) was used to consider birds with both migratory statuses. Wintering birds with spring and autumn migratory populations were taken into consideration with an overall November-February trapping effort, while birds mostly in breeding status but with spring and autumn migrant populations were taken into consideration with an overall March-October trapping effort. The annual and seasonal abundances and trapping effort by species are presented in Table 1. According to Poulin et al. (2000), the normalized breeding trapping effort used to measure the assemblage of reed bed species was set at 1.300 m2/hour, which satisfies the requirements at least during the study site’s breeding seasons.

**Table 1.**
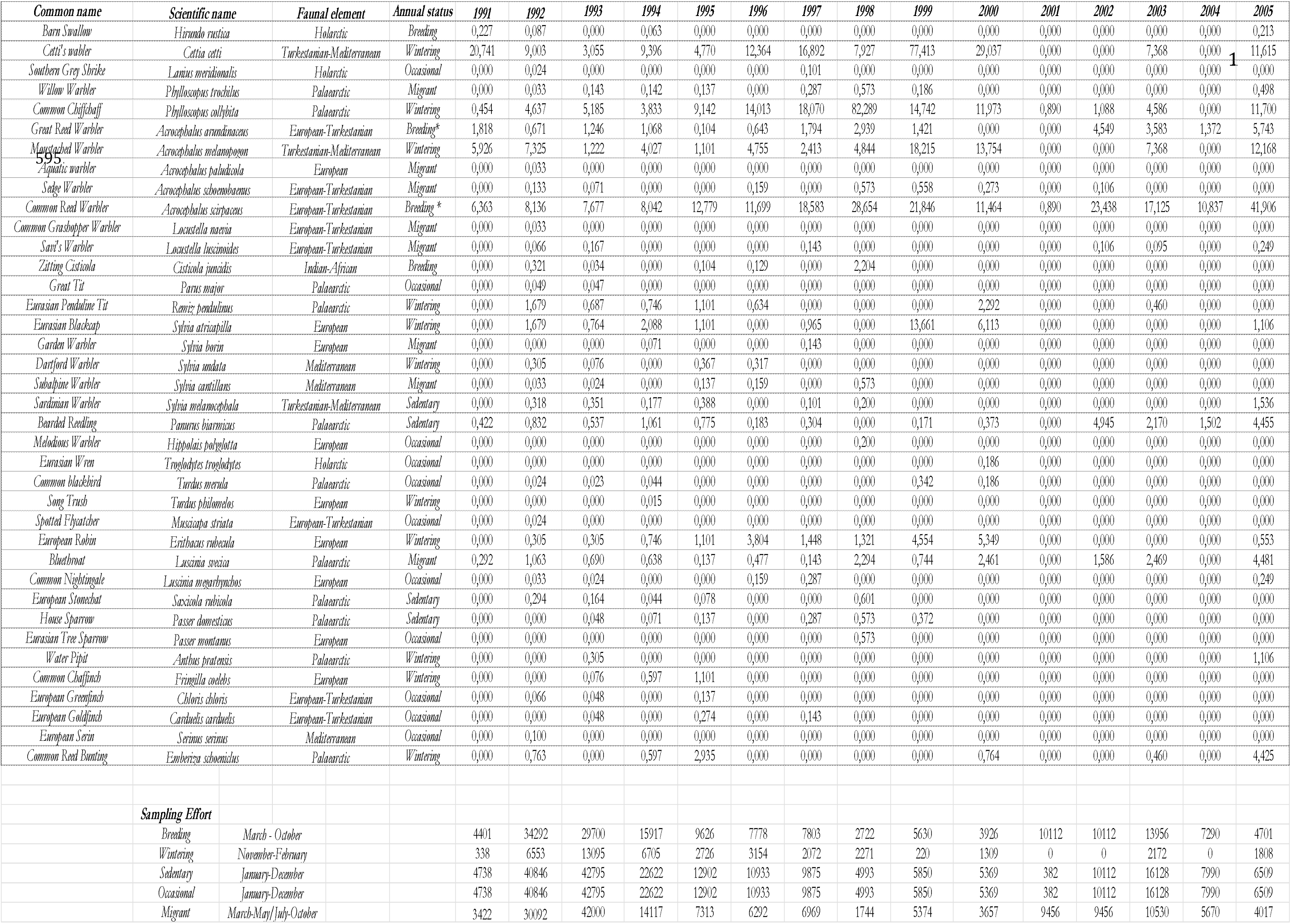
List of species ringed in EH in 1991-2005. Standardized abundances of each species in mist nets [individuals/m2net* hour) *1000] are plotted. Abundances were calculated differently according to the annual occurrence of the trapping effort at the site. Breeding (March-October, B), wintering (November-February; W), sedentary (January-December, S), spring migrant (March-May, SM), autumn migrant (July-October, AM), occasional (O). Birds with both migratory phenology were considered as an adding spring-autumn trapping effort. Birds with a mainly breeding status but with spring and autumn migrant populations, marked as (*), were considered with an overall trapping effort March-October and wintering birds with spring and autumn migrant populations, marked as (*) were considered with a summed trapping effort November-February and June-September. Summed total abundances between 1991 and 2005 are also plotted for each species. Abundances were transformed into indices for TRIM analyses. Birds were classified and English named according to Rouco et al (2022). Faunal elements follow Voous (1960).

I performed a checklist of passerines from EH that were initially categorized into faunistic elements by Voous (1960) and listed, classified, and given English names by Rouco et al. (2022). The plotting of this general template was the same in Tables 1-4. I calculated the individual biodiversity distributions (Shannon’s Index) between periods (D1 and D3) to see whether or not the bird assemblage was similarly distributed between periods (Van Turnhout et al 2007). This allowed me to determine whether the bird assemblage may have altered between then. Since bird diversity between assemblages did not meet normality (Shapiro Wilk’s tests: D1 = 0.585; P < 0.001; D3 = 0.008; p < 0.001), median diversities were evaluated using the non-parametric Mann-Wittney-Wilcoxon rank sum tests (MWW, Sokal & Rolhf 1981). Because the relative number of specialists and generalist species is likely to affect community functions (Julliard et al. 2006), I attempt to provide a suitable index of specialisation of each species within the assemblage using the species-specific specialisation index (SSI) derived from Morelli (Appendix 1 in Morelli et al. 2020). According to DeVictor et al. (2008b), this index is based on how frequently each species occurs in each type of habitat or land use. As a result, each bird species’ specialization index (SSI) may have values ranging from 0 to 1, which correspond to low and high levels of specialization, respectively. The specific weighted SSI of the assemblage in each time was calculated by multiplying the SSI index for each species by the number of individuals. The abundance of each species, as indicated by the number of individuals, is taken into consideration when computing the ensemble of community specialization in this index. In order to determine whether the two periods were equally distributed on time and because the SSI between assemblages in the period D1 and D3 did not meet normality (SSI-1 = 0.585; P < 0.001; SSI-3 = 0.008; p < 0.001), the median weighted SSI for D1 and D2 were compared using the Mann Wittney Wilcoxon tests for two independent samples.

### 2.5. The Population trends and climatic and management effects

For analyzing species-specific trends, I employed Trends & Indexes for Monitoring TRIM software (Pannekoek & Van-Strien 2001), which was developed from R-Project v. 44.2 software (R-Core Team 2020) by the “rtrim” package (Bogaart et al. 2024). I implemented “rtrim” using Model 2 based in Poisson regression method (Van-Strien et al. 2004, Pannekoek & Van-Strien 2001) which accounts for only one slope in the overall model (Ter Braak et al. 1994). To indexing the bird abundances data and run the TRIM they were transformed into integers. Null zero assignments were one-added and missing values were transformed into not analyzed NA values and afterwards imputed (Ter Braak et al. 1994). See Table 2 for complete explanations. Using the “glm2” package (Marschner 2011), I employed *Generalized Linear Mixed Models* (GLMM; McCullagh & Nelder 1983) to find some relationships between climate and habitat transformation. The response variable was the abundance of each bird, and the predictive variables were annual mean temperatures, annual total precipitation, and reed height. For more details on the analysis, refer to Table 3.

**Table 2.**
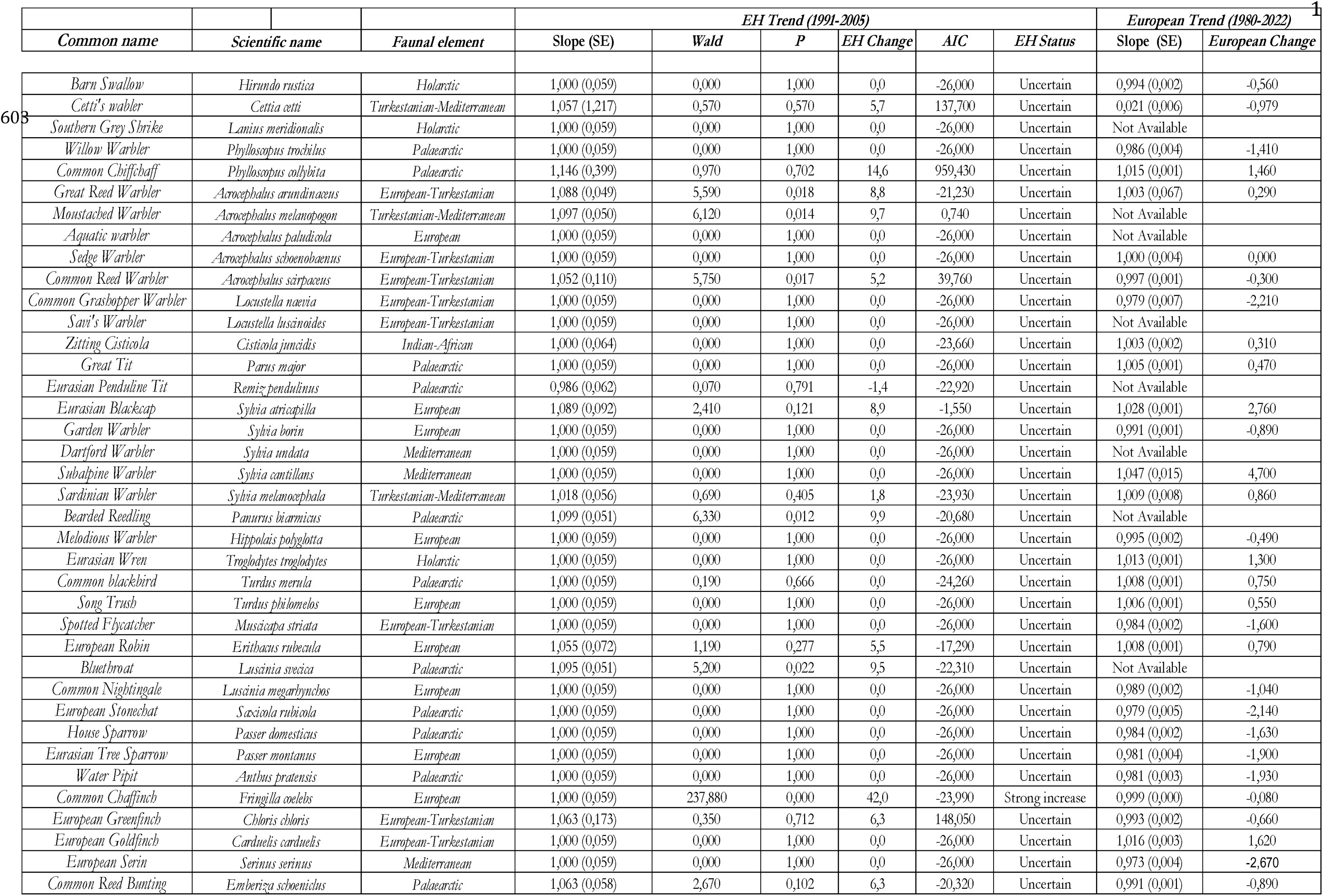
Species trends in EH for D1 period in EH and in Europa (1980-2022) (2) based on implemented TRIM software (Pannekoek & Van-Strien 2001). The overall trend based on multiplicative slopes jointly with Standard Errors (SE) are depicted. If the slope value is 1, there is no trend. If >1, there is a positive trend, if <1, the trend is negative. The percentage of change from imputed data is estimated from the slope. For instance, 1.08 means an 8% increase per year, 0.93 means a 7% decline per year. Data overdispersion was tested by the Wald test (values > 1 and significant and indicates that there is overdispersion and they fit a Poisson Regression equation. Trend status of each species is as the pointed out by Vorisek et al (2008). Best models are less sparse and those with greater Akaike’s value (AIC). Data from (2) was obtained online in https://pecbms.info/trends-and-indicators/species-trends/.

**Table 3.**
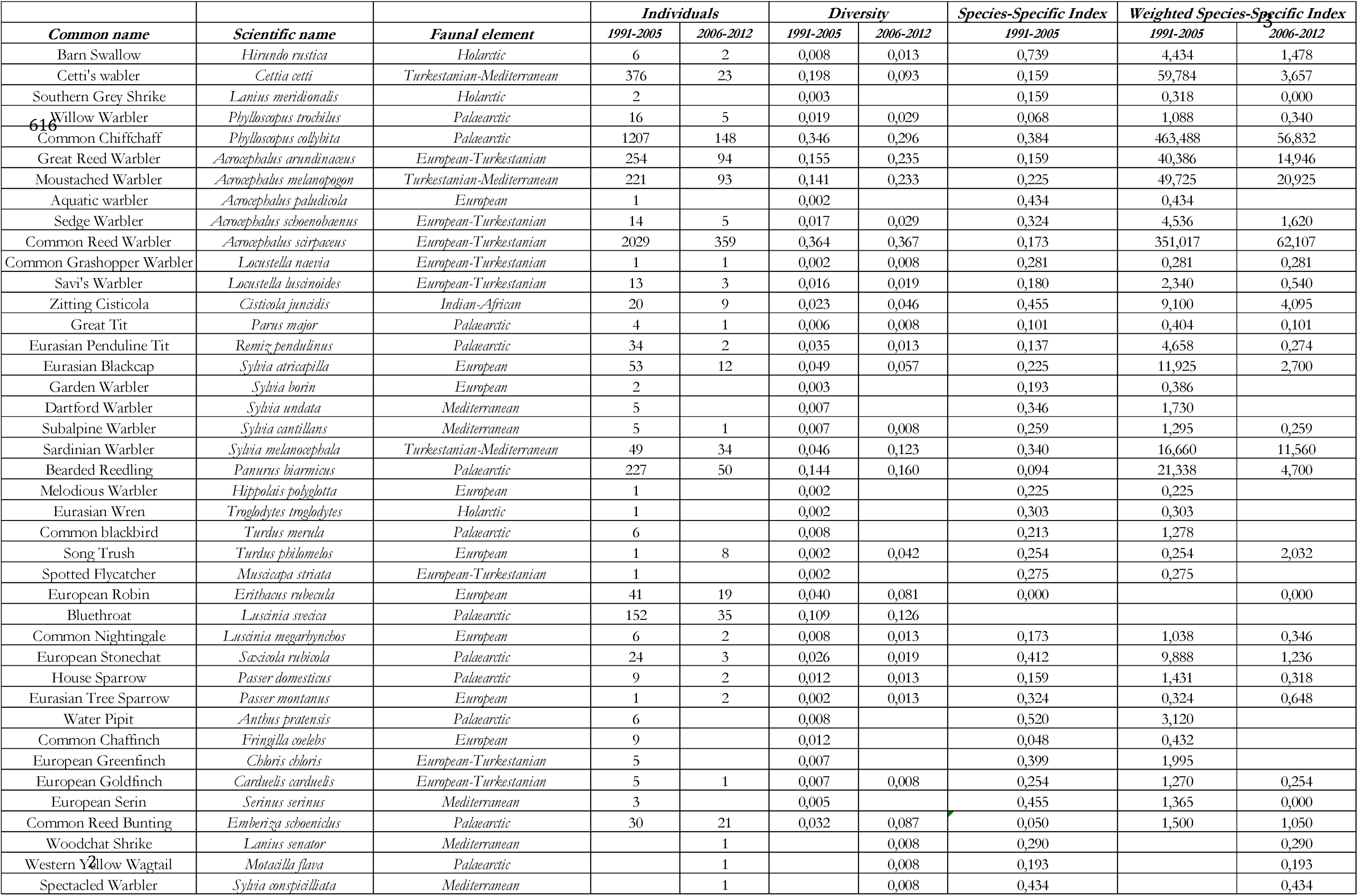
General diversity classification based on Shannon’s Diversity Index for D1 and D2 periods. Species Specific Index (SSI-1 and SSI-2, see methods) are depicted only for D1 period.

## 3. Results

### 3.1. The overall climatic sketch and trends of weather variables

The average temperature during the coldest month (January, period D2) was 19.4 ± 1.54, while the average rainfall during the warmest month (July) was 1.4 ± 2.99 mm. The total annual precipitation averaged at 241.8 ± 88.14 mm, with an average annual temperature of 19.4° ± 0.96 recorded for the period of 1991-2020. Over the period D1 annual temperatures in EH decreased (Temperature = - 0.029·Year + 76.399; r^2^ = 0.018, F_1,13_ = 0.2372, P = 0.634) and rainfall increased (Precipitation = 1.920·Year – 3610.731, r^2^ = 0.016, F_1,13_ = 0.219, P = 0.648) these trends were similar in D2 (Temperature: Pearson’s r = - 0.077, P = 0.685; Rainfall: r = 0.286; P = 0.126). Large scale temperatures were influenced spring temperatures since they got higher determination coefficient on the linear relationships (r^2^ = 0.723, P < 0.001). Likewise, rainfall in current autumn was the greatest driver of overall precipitation (r^2^ = 0.521, P < 0.001). Punctual storminess may lead extreme rainfalls as occur in other central Mediterranean areas (Diodato & Bellocchi 2010) and determinates the highest variability of rainfall (CV = 35%) compared to temperatures (CV = 5%) since outliers on the large-scale precipitations in 2019 (469 mm; Grubbs Tests, G = 2.509, U = 0.776, P = 0.124) could be responsible for the positive correlations of rainfall with year so this scenario represents an apparent dry and medium-warm cycle in the study area during the last century (1991-2020) which could extend to the southern subregion of Alicante and subsequently, due to climate change, rainfalls became shorter but more intense in an episodic manner (Diodato et al 2011) so in climatic terms, conformed a posterior cycle (1921-1924) in a wetter and warmer period. This is coincident with the great local climate variability in Mediterranean areas depicted by Xoplaki (2002).

### 3.2. The Zoogeography and trends in the avifauna

Seventy percent of the assemblage’s species (N = 38) were of Palearctic, European, and European-Turkestanian origin. A reminiscence was of Turkestanian-Mediterranean origin. An annual trapping effort (r = 0.800, P < 0.001) that grew weakly on time (r = 0.073, P = 0.795) had a considerable influence on the yearly total number of species, which declined weakly on time (r = - 0.092, P = 0.734). The sole species with European status, the Common Chaffinch *Fringilla coelebs*, saw a significant rise (42%) over the 1991–2005 study period, while the Eurasian Penduline tit *Remiz pendulinus* saw a reduction of 1.4%.

During the study period, the remaining assemblage had an ambiguous trend status with a positive trend slope ranging from 0 to 15%, primarily non-significantly. Even though the overall diversity was 13% greater in the following years (1991-2005: 1.875 nats, 2006-2012: 2.163 nats), the median diversity across periods was statistically significant (D1: 0.010 nats, D3: 0.029 nats; Wilcoxon-Mann-Wittney Test; W = 351.5, P = 0.012). Additionally, the community became increasingly less specialized over time as a result of the overall SSI index declining by 82% (D1: 1070.3, D3: 193.2) as a result of 32% of species (N = 12) being lost and three species being added: two Mediterranean, the Woodchat shrike *Lanius senator* and the Spectacled warbler *Sylvia conspicillata* and one Palearctic, the Western yellow wagtail *Motacilla flava* (Table 3).

### 3.3. The relationships with climate and habitat transformation

Simple (linear) models that primarily used one variable to explain annual abundance did not significantly fitted most species (Table 4). 78% of the species in the collection were influenced by climate factors, particularly temperature and precipitation, which mostly determined the species’ interannual abundance. Most of these species were Palearctic (16%) and Euro-Turquestanian (21%). 29% of species were impacted by habitat change, primarily in the Palearctic (13%).

**Table 4.**
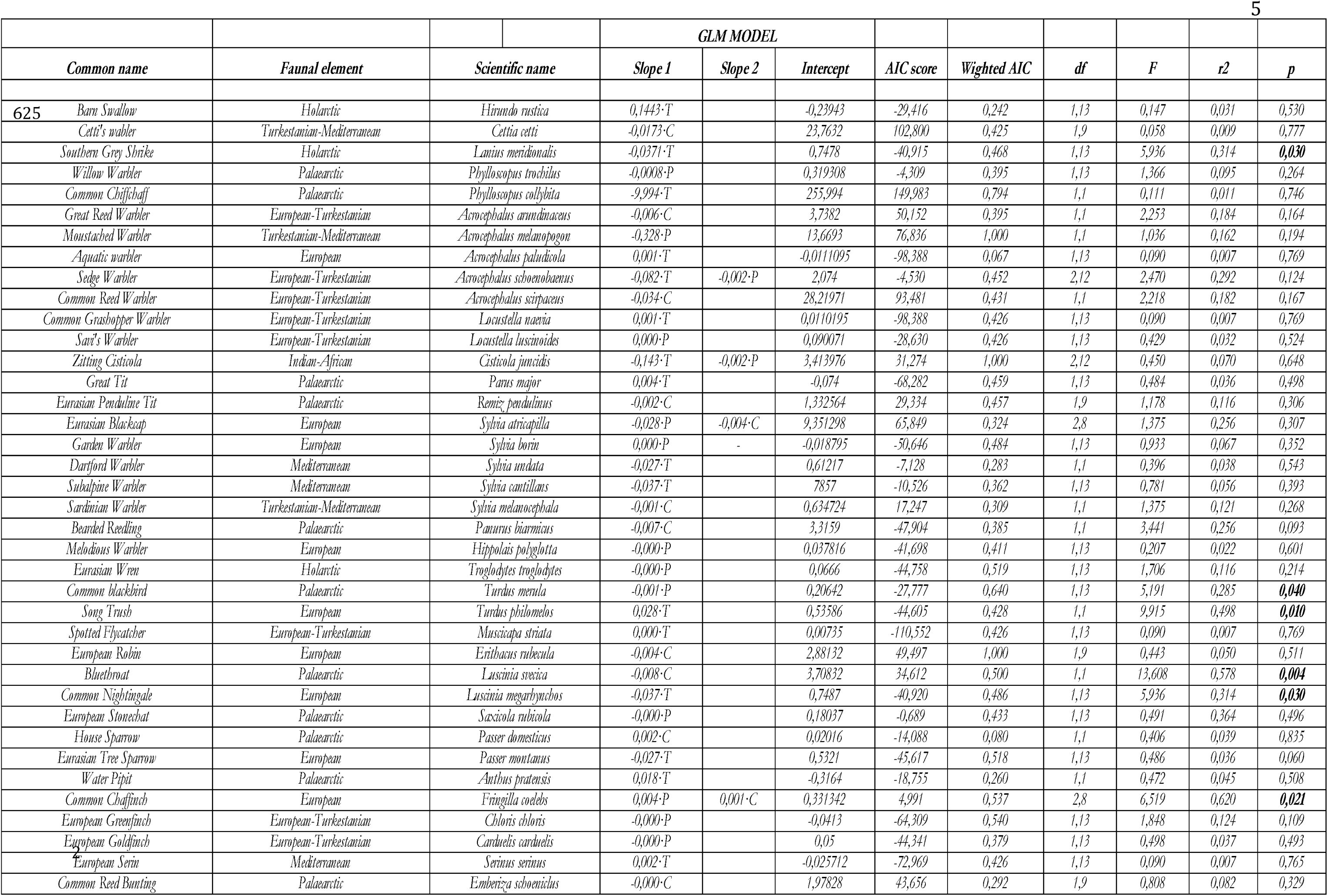
General Linearized Mixed Models (GLMM) implemented by package “*glm2*” (Marschner 2011) in the R-Project software (R-Core Team 2921) on the specific abundances as fixed dependent variables with climatic and management variables as response-proxy independent variables. Only the main interactions among independent variables were considered. Slopes of the Linear Regression Models are depicted. Minor AIC’s values were used to select the best model. ANOVA’s F-value with degrees of freedom and determination coefficient (r2) of the selected model and p-values of the regression. Significant (P < 0.05) in bold. C = habitat; T = Temperature; P = Precipitation

## 4. Discussion

Southeastern Iberia experiences a Mediterranean climate characterized by hot, dry summers and mild, wet winters. The primary meteorological factors influencing this climate include: 1.-Temperature variability: rising temperatures have been observed in the Mediterranean region, significantly affecting local ecosystems and species distributions. For instance, the abundance of migratory birds like Savi’s warbler *Locustella luscinioides* has shown a positive correlation with intra-annual temperatures, indicating that temperature fluctuations can impact breeding and migratory patterns (Peiró 2024), 2.-Precipitation patterns: precipitation in SE Iberia is characterized by seasonal variability, with significant impacts on local flora and fauna. The region has experienced decreasing precipitation trends, which can lead to drought conditions, particularly in summer. This has been linked to the adaptation and migration of various species, as well as changes in vegetation dynamics (Carrasco et al. 2018, Alambiaga et al. 2021) and 3.-Climatic oscillations: the climate dynamics in SE Iberia are also influenced by larger climatic patterns such as the North Atlantic Oscillation (NAO) and the El Niño Southern Oscillation (ENSO). These oscillations can lead to variations in precipitation and temperature, affecting the overall climate stability in the region. These factors collectively shape the unique climatic conditions of southeastern Iberia, influencing biodiversity, species distribution, and ecological interactions.

The findings indicate that the majority of the species in EH are generalists, with only a limited number of reed-marsh specialists (*Acrocephalus*, *Cettia*, *Locustella*, Cramp 1992, Kennerley & Pearson 2010) making up 25% of the bird assemblage. The arrival of four species of passerines, not specialists, was the primary cause of the diversity increaseThe Zoogeography of EH is primarily Palearctic, European and European-Turkestan, owing the wintering of Palearctic wintering heavyweights of Common Chiffchaff *Phylloscopus collybita* (PC) and the summer European-Turkestan Eurasian Reed Warbler *Acrocephalus scirpaceus* (AS). These two (AS and PC) are quantitatively the bulk species, increasing in abundance at rates of 5–14% each year, respectively. However, they are adversely affected by habitat fragmentation and annual temperature increases. There may be a decline in the PC population but a rise in AS in the future because to rising temperatures and habitat resiliency.

In the 1960s, Turkestan-Mediterranean elements, such as scrub warblers like the Sardinian Warbler *Sylvia melanocephala* (SM) and two reed-marsh specialists, the Cetti’s Warbler Cettia cetti (CC) and the Moustached warbler *Acrocephalus melanopogon* (AM), spread towards central Europe from southern Europe and southwest Asia. More recently, in the case of AM, they likely came from wintering birds of populations of central and eastern European origin (Bozó et al. 2023), and there may have been an over-dispersal of breeding populations from the eastern Iberian regions to the south (Ceresa et al. 2015).

Diversity in 1991-2005 in EH falls well below the values detected in other Mediterranean wetlands (Malavasi et al. 2009), indicating that the study area is human-disturbed and is highly below the range of diversity of natural or semi-natural central Mediterranean wetlands (2.8-2.5 nats, Malavasi et al. 2009). The huge increase in diversity from 2006 to 2012 is indicative of transformation towards a natural wetland.

Denser reeds and less rainfall appear to be more conducive to the population dynamics of the earlier specialists, whose abundances were adversely affected by habitat management and rainfall, respectively. We could anticipate a notable rise in the population of Turkestan-Mediterranean elements (AM & CC) in this ecosystem in the event of future warming, less precipitation, and the restoration of the natural habitat in EH. Since reedbed alterations (clearing) have a detrimental impact on SM, boosting habitat densification would have a significant favourable impact on their abundances.

The Reed bunting, another reed marsh specialist, is only found in EH as an overwintering bird and has not been breeding there since the 1970-80s (Medina & Garcia 1984, Medina 1985). It is likely that the earliest breeders here were of the Iberian subspecies *witherby*, which is found in the widely recognised breeding nucleus in the central (Alambiaga et al. 2021) and eastern marshlands of Iberia (Carrasco et al. 2018).Simple (linear) models that explained the annual abundances on climate and habitat alteration were mostly not significant and this is in accordance with the findings of Jiguet et al (2010) which hypothesizes that trends in European abundances do not meet lineal relationships. According to Leviol et al (20102), European bird communities are dominated by native generalist species, indicating a significant homogenization process in different regions. The authors propose that this rapid and non-random change in community composition is due to anthropogenic activities, but the effects of climate change are also very important (Davey et al 2011).

## Conclussions

Future research during longer cycles is needed to ascertain the possible changes in the species’ composition due to climate warming and management use. To determine the potential changes in the species’ composition brought on by climate change and management use, more research is required.

## Notes

### Competing Interest Statement

The authors have declared no competing interest.

### Summary of Updates

The Mediterranean region is facing significant impacts on populations of biota, particularly birds, due to climate variability and habitat alteration. Consequently, the ecological diversity of many ecosystems has diminished, and the abundance of bird species has declined. This study employs a novel strategy in a seasonal freshwater artificial wetland in south-east Iberia establishing species-specific index trends for common passerine birds by comparing them with climate and habitat fragmentation data. A climatic overview indicates minimal trends towards warmer and wetter conditions in terms of temperature and rainfall over the 15-year study period. This pattern persisted over a 30-year period, resulting in the formation of a Mediterranean climatic subregional scenario with warm, dry conditions that may influence the passerine assemblage in an analogous manner to that observed in other Mediterranean assemblages. One notable feature is that this assemblage in the 1990s was characterised by low natural diversity and specialisation when it was a highly disturbed wetland. However, it became natural by the mid-2000s, with diversity values like those observed in other central Mediterranean wetlands. Trends used to evaluate species-specific patterns revealed that 98% of species had an unknown trend status during the study period, with a trend slope ranging from 0 to 15%, which was not substantially different than zero, probably due to the unequal standardisation methodology and the short time studied. Further research using Generalized Linear Mixed Models (GLMM) revealed that precipitation and habitat alteration had the most detrimental effect on species. Of the observed species, 60% were affected by both factors, the majority of which were Palearctic (20%). The reduction of wintering areas and shifting migration routes may account for these effects. Further research is required on longer temporal data to clarify the evolution of the avifauna in this wetland.

